# Seed origin determines cork oak germination: the warmer the higher, faster and more synchronized

**DOI:** 10.1101/2025.10.03.680296

**Authors:** Marion Carme, Eduardo Vicente, Filipe Costa e Silva, Maurizio Marchi, Giovanni Giuseppe Vendramin, Natalia Vizcaíno-Palomar, Boutheina Stiti, Marta Benito Garzón

## Abstract

The early life stages of trees, particularly germination, are crucial to fitness and highly sensitive to climate. The influence of temperature on recalcitrant seed germination has rarely been studied due to their desiccation sensitivity, which hampers storage. However, Mediterranean recalcitrant oaks would be particularly affected by the expected increased temperature in this region. Here we investigated the effect of warming temperatures on germination of 975 acorns from 8 range-wide *Quercus suber* populations. We sowed the acorns at 15, 20 and 25 °C in climatic chambers, and monitored germination during 4 months. The germination dynamics in each chamber was explored by a Cox proportional hazards model. We assessed environment (germination experiment temperatures), population (climate of seed origin) and their interaction effects on germination percentage, time, and synchrony using generalized linear mixed-effects models. Genetic clines on germination percentage, time and synchrony were mostly triggered by temperature, with seeds from warmer origins showing higher germination, earlier timing, and greater synchrony than colder ones. Higher sowing temperatures promoted advanced germination, and this effect was higher in seeds originating from regions with stronger seasonality. Earlier and synchronous germination found in seeds from warm origin may reduce the desiccation probability for acorns and seedlings, while late germination and low synchrony found in seeds from cold origin might be an adaptive response to unpredictable frost events that would impair seedling survival. The germination synchrony adaptive response was unexpected and further investigation on recalcitrant seeds’ germination dynamics in response to increased temperatures is needed to confirm it.

## 1. Introduction

Climate change is increasing the frequency and intensity of drought, fire, storms and pest outbreaks in forests worldwide (Menezes-Silva et al., 2019). As a consequence, tree populations can extirpate, migrate or survive *in-situ* by adjusting to new conditions through evolutionary and ecological strategies (Aitken et al., 2008). These responses along large environmental clines are notably understudied in early life stages in comparison with adult ones (Alberto et al., 2013; Leites and Benito Garzón, 2023), despite them being crucial to survival and reproduction (Verdú and Traveset, 2005), key steps to adapt to new environmental conditions (Donohue et al., 2010; König et al., 2022) and more sensitive to climate than adult trees (Erlichman et al., 2024; Ettinger and HilleRisLambers, 2013).

Among early life stages, germination has notable impact on the entire life cycle of trees through cascading effects (Donohue et al., 2010; Gremer et al., 2020). It is a fitness component which determines how many individuals will reach the seedling stage (Donohue et al., 2010). The phenology of germination determines the environment under which the seedling grows and is tuned to reduce environmental risks, e.g. avoiding late frost in temperate species (Donohue et al., 2010; McCartan et al., 2015) or summer drought in Mediterranean ones (Cruz-Tejada et al., 2024). Further, adaptive processes on population germination dynamics can help seedlings to survive under unpredictable environments. For instance, species forming seed banks (i.e. orthodox dormant seeds) growing in unpredictable environments typically show bet-hedging strategies involving low synchrony in germination that benefits the long-term population growth rate (Abley et al., 2024; Gremer et al., 2016; Maleki et al., 2023). On the contrary, recalcitrant (i.e. desiccation sensitive) non-dormant seeds are expected to show high germination synchrony to avoid desiccation after shedding (Maleki et al., 2023).

Temperature is a key factor controlling seed germination traits (Roberts, 1988) and population-level differences are often linked to temperature at seed origin (Chamorro et al., 2018; De Frenne et al., 2012; Zettlemoyer et al., 2017). In addition, germination traits typically exhibit a high plastic response to temperature fluctuations. In general, warming accelerates the metabolic reactions that activate germination (Ali et al., 2017), which tends to increase germination percentage and to hasten germination time (Fernández-Pascual et al., 2021; Milbau et al., 2009). Nonetheless, the opposite effects can be observed if warming exceeds the germination temperatures at which seeds are adapted (Maleki et al., 2024; Sentinella et al., 2020). Hence, the effects of temperature can differ greatly among populations of the same species, particularly in broad-range ones. For instance, contrasting germination responses to warming have been documented between northern and southern populations of *Acer saccharum* (McCarragher et al., 2011; Solarik et al., 2016) and *Betula pendula* (Solé-Medina et al., 2020). However, most range-wide studies about temperature effects on germination traits are based on species with orthodox seeds (Fernández-Pascual et al., 2021; Milbau et al., 2009), as recalcitrant seeds cannot be stored and they can quickly lose viability during transport (Berjak and Pammenter, 2008; Wyse et al., 2018). Yet, many keystone tree species, particularly in Mediterranean climates, produce recalcitrant seeds (Tweddle et al., 2003). Thus, in a context of climate change, it is critical to address the knowledge gap about temperature effects on the germination traits of recalcitrant species.

Cork oak (*Quercus suber* L.) is a drought-tolerant Mediterranean species with recalcitrant seeds (Aronson et al., 2012), that can adapt to contrasting climate regimes by adjusting key fitness-related traits (Benito Garzón et al., 2024; Ramírez-Valiente et al., 2022, 2015, 2014; Vicente et al., 2025). It mostly germinates in autumn and in spring (Aronson et al., 2012), as these periods generally provide enough moisture and mild temperatures in Mediterranean climates, especially in coastal areas (Harding and Palutikof, 2009). Germination rates are high, particularly in seeds from drier origin. In a recent study, Benito Garzón et al., (2024) found population differentiation in germination time and probability, with warmer populations being prone to an earlier and higher germination. However, this study was performed with partially desiccated acorns, which can notably alter germinability and germination time (Amimi et al., 2023). Thus, the isolated effect of temperature variation and how it is modulated by populations’ climate remains poorly understood. Moreover, most germination studies focus on germination percentage or time, while germination synchrony has been overlooked.

We investigated the effect of rising temperatures on germination percentage, time and synchrony across range-wide cork oak populations. To this aim, we conducted a germination experiment using acorns from eight populations under controlled conditions at 15°, 20° and 25°C. Water availability was maintained constant during the experiment to isolate the effects of temperature. We analyzed germination percentage, time and synchrony using Cox proportional hazards models, and generalized linear mixed-effects models. Our objectives were to: (i) identify potential genetic clines to temperature in germination percentage, time and synchrony along the climatic gradient occupied by the species; (ii) assess the effect of warming temperature during the germination process; (iii) investigate how germination plasticity to temperature varies among populations, i.e. the interaction between environment and population.

## 2. Material and Methods

### 2.1. Plant material

We collected acorns from eight natural populations distributed across the species distribution range in Italy, Tunisia, Portugal, Spain, and France (Fig. 1) during October, November and December 2022. “Population” was defined as a group of at least 10 mature individuals within a single forest stand and thus experienced the same local environmental conditions. In each population, we sampled 6–10 mother trees (65 in total) and collected ~25 acorns per tree to assure 15 acorns for sowing (i.e. 90-150 acorns per population). This sample size was appropriate for our objective of capturing population-level responses while accounting for within-tree variance (Mérian et al., 2013), especially since *Quercus* species typically show high germination percentages (Bonner & Karrfalt, 2008). We only collected those acorns that dropped after shaken the tree branches to assure the same maturation status for all acorns (i.e. brown pericarp and without radicle emergence). The geographical coordinates of the populations, mother tree circumference (used as a proxy of maternal effect) and collection date were recorded for all sampled trees (Fig. 1). Before sowing, we imbibed the acorns in water for 10 minutes and removed the floating ones to sort out the healthy acorns. In total, we used 965 acorns.

**Fig. 1:**
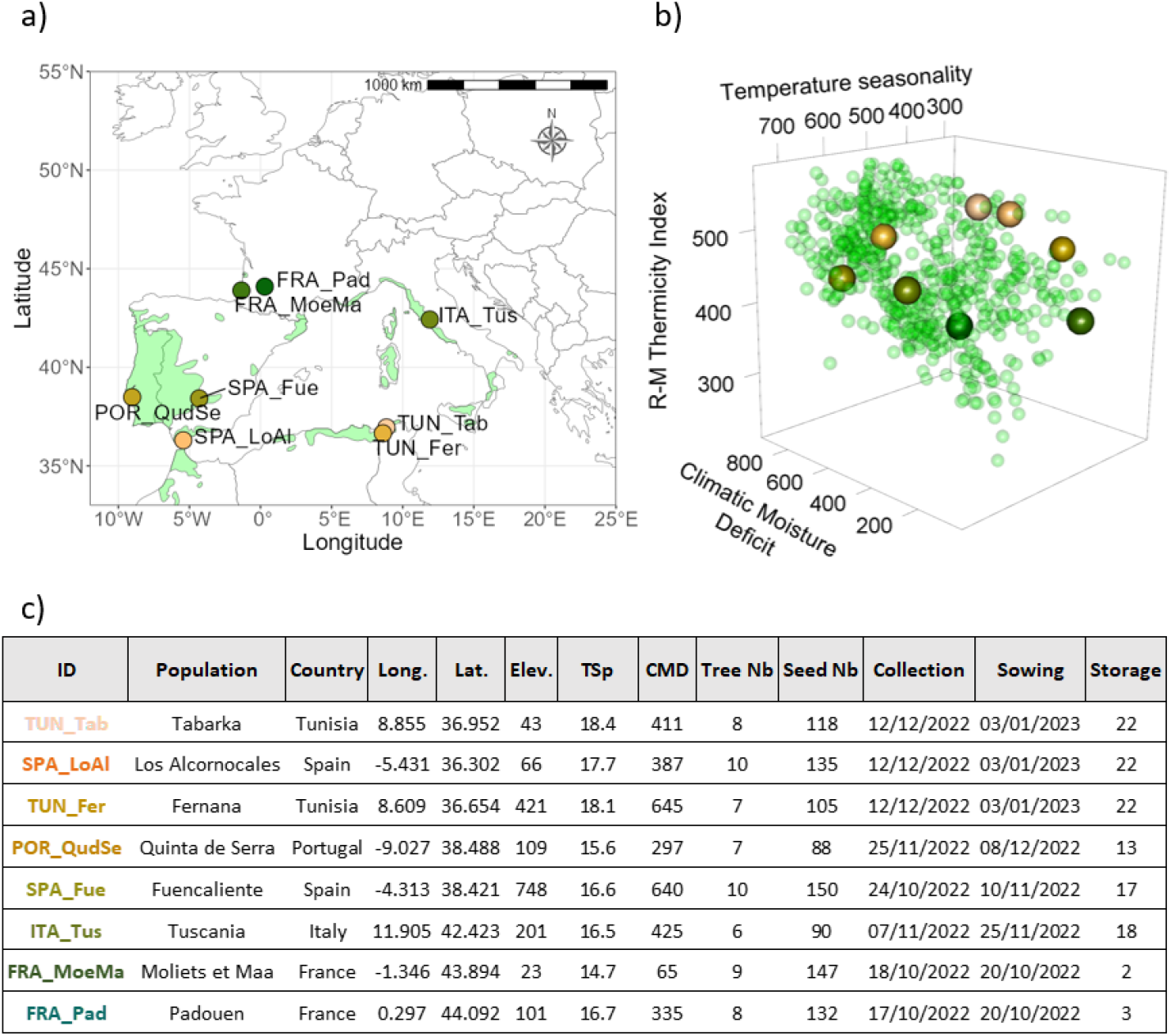
Cork oak acorn populations within the species distribution range (EUFORGEN, (Caudullo et al., 2017) in a), within the species’ climatic envelope according to Climatic Moisture Index, Rivas-Martínez Thermicity Index and Temperature Seasonality in b) and details in c), with ID sorted by Thermicity Index; Elev. = Elevation (m); TSp = maximum spring temperature (°C); CMD = Hargreaves Climatic Moisture Deficit (mm); Tree Nb = number of trees per population; Seed Nb = number of acorns per population. Populations colors indicate their origin temperatures (warm colors for warmer populations and cold colors for colder populations, according to Rivas-Martínez Thermicity Index).

### 2.2. Germination experimental design

Acorns were sown right after collection to avoid desiccation; their sowing date varied according to each population’s collection date. Hence, the experiment started on October 2022, and lasted until the beginning of February 2023, assuring that germination was monitored for 90 days for every acorn. Acorns were put in nursery trays (4×5 pots), in 6×6×8 cm pots (one acorn per pot), filled with potting soil (NPK 8-2-7, dry organic matter 80%, conductivity 30mS/m, pH 6.5, water retention 820ml/L), with the apical end exposed at the soil surface, allowing direct observation of radicle protrusion.

To test the temperature effect on germination, we sowed the acorns in three climatic chambers (Snijder LABS, micro clima-series) at 15, 20 and 25 °C during the day, and 10, 15 and 20 °C during the night (325 acorns per chamber). These temperature regimes approximate the temperature range at seed origin during the germination period (Table S1) and include a +5 °C increment to simulate projected warming under the SSP5-8.5 scenario. Other environmental parameters were kept constant: 75% relative air humidity, 300 μmol.m-2.s-1 light intensity (Li-190R Quantum Sensor), photoperiod of 13/11 light/dark hours, and watering twice a week with distilled water.

### 2.3. Germination measurements

We measured individual acorn mass before sowing (precision of 0.1 g). Germination monitoring was carried out thrice a week. An acorn was considered as germinated when radicle had emerged by 2 mm. We measured individual germination time (T0 = the number of days between sowing and emergence) and germination status (G = 1 if germinated, 0 if not).

From T0 and G, we calculated three indexes with the *germinationmetrics* 0.1.8 R package (Aravind et al., 2023) for each combination of population – chamber – mother tree:

Averaged germination time (AT0 = the mean number of days between sowing and emergence), calculated as:

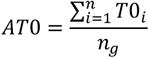

where *T*0_*i*_ is the individual germination time of each germinated acorn, and *n*_*g*_ the total number of germinated acorns within the group

Germination percentage (GP = number of germinated acorns at the end of the experiment), calculated as:

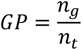

where is the number of germinated acorns and the total number of sown acorns within the group Germination synchrony (GS = spread of germination over time), calculated with the following formula (Ranal and Santana, 2006):

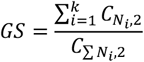

where 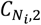 is the number of pairwise combinations among the *N*_*i*_ seeds that germinated in the *i*-th time interval (estimated as 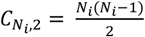), and 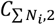 is the total number of pairwise combinations among all seeds that eventually germinated, assuming that all germinations occurred simultaneously.

G - GP, T0 - AT0, and GS represent the three main aspects that characterize germination: germinability, speed and spread over time, respectively (McNair et al., 2012; Ranal and Santana, 2006).

### 2.4. Climatic characterization of populations

We characterized the populations according to their climate of origin. We particularly aimed to investigate the long-term effects of climate on germination, based on the rationale that historical climatic conditions have shaped the adaptation of populations to climate. To this end, we retrieved 28 climatic variables from ClimateDT over the 1901-1960 period (Marchi et al., 2024), i.e the earliest available data to 1960, which we use as the proxy of the current climate change onset (Fichefet and Tricot, 1992; Garcia-Mozo et al., 2015).

We conducted a PCA to select non-correlated climatic population variables (Fig. S1). We preferentially selected indices that are more integrative of various climatic components. We ended with the variables reported in table 1: *CMD* the Climatic Moisture Deficit (population water availability); *RMTh* the Rivas-Martínez Thermicity Index (population temperature); *TSp* the maximum spring temperature; *TSn* the temperature seasonality (inter-annual temperature variability).

**Table 1:** Population climatic variables selected to characterize populations and investigate population effect on germination traits in generalized linear models.

| Name and Acronym | Unit | Description | Calculation |
| --- | --- | --- | --- |
| **Hargreaves Climatic Moisture Deficit (CMD)** | mm | Considers water stress in plants | Hargreaves reference evapotranspiration (computed with the SPEI R package) minus mean annual precipitation (Hargreaves and Samani, 1985) |
| **Rivas-Martínez Thermicity Index (RMTh)** | °C | Considers the intensity of the cold (limiting factor for many plants), together with the average annual temperature | Sum of average annual temperature, average temperature of the minimums of the coldest month, and average temperature of the maximums of the coldest monthly period (Rivas-Martínez et al., 2002) |
| **Temperature seasonality (TSn)** | °C | Temperature change over the course of the year | Standard deviation of mean monthly temperatures |
| **Maximum spring temperature (TSp)** | °C | Considers the spring temperature where germination occur | Mean of maximum temperatures in March, April and May |

**Table 2:**
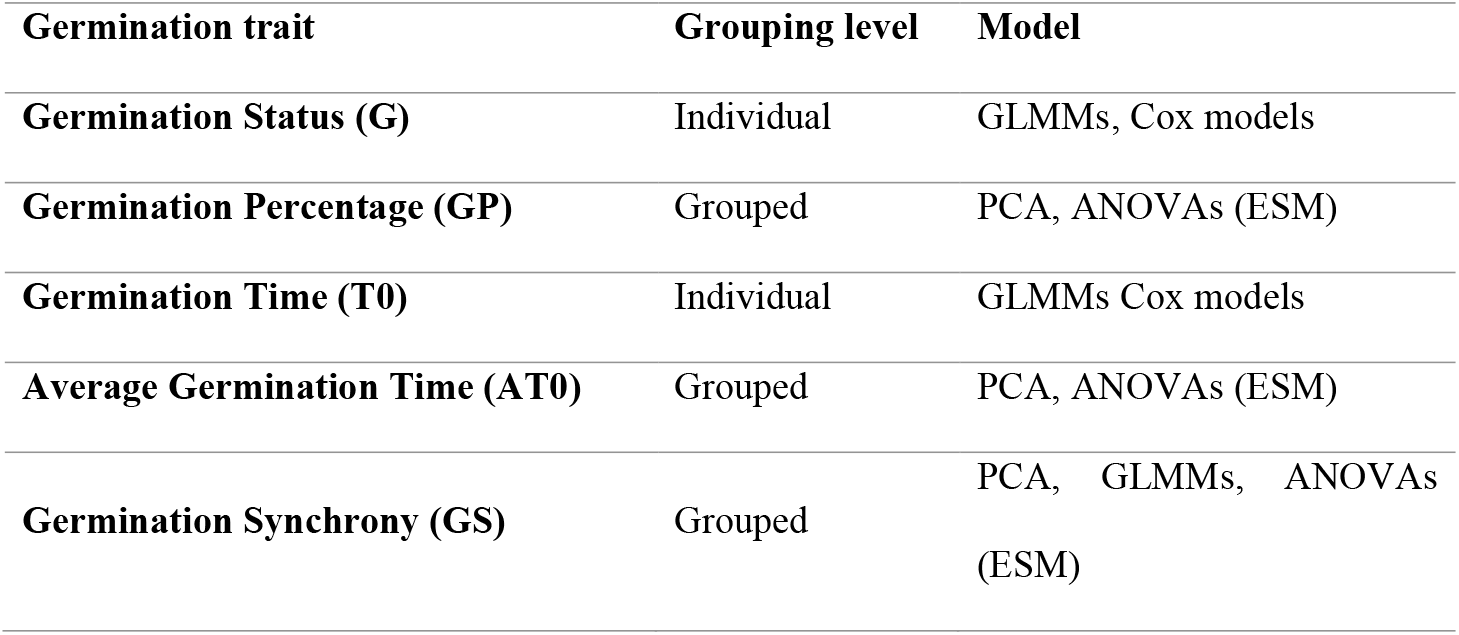
Germination traits considered and their analysis. Germination traits = germination traits observed and derived indexes; Grouping level = whether the trait was measured at the individual level or aggregated per population–chamber–mother tree; Model = type of model used for each trait. Because we aimed to maximize statistical power by working at the individual level whenever possible, we analyzed germination status (G) and germination time (T0) using mixed-effects models. Germination synchrony (GS), by definition a group-level index, was analyzed at the population–chamber–mother tree level. We also calculated germination percentage (GP) and average germination time (AT0) to allow the analysis of germinability and germination time in PCA, ANOVAs and pairwise comparisons, and to check the robustness of covariate effects observed in the G and T0 mixed-effects models.

| Germination trait | Grouping level | Model |
| --- | --- | --- |
| Germination Status (G) | Individual | GLMMs, Cox models |
| Germination Percentage (GP) | Grouped | PCA, ANOVAs (ESM) |
| Germination Time (T0) | Individual | GLMMs Cox models |
| Average Germination Time (AT0) | Grouped | PCA, ANOVAs (ESM) |
| Germination Synchrony (GS) | Grouped | PCA, GLMMs, ANOVAs (ESM) |

### 2.5. Statistical analysis

We first conducted a time-to-event analysis to understand the overall germination dynamics and compare hazard rates across growing chambers and populations. Then, we performed different mixed-effects models to analyze the germination status (G), time (T0) and synchrony (GS) of germination. All analyses were carried out in R version 4.3.2 (R Core Team, 2023) and script is available at *available upon publication*.

#### 2.5.1. Time-to-event analysis: Cox proportional hazards models

To estimate the differences in germination probability over time according to population and experiment temperature, we used Cox proportional hazards models (Cox, 1972). They consider the entire time-course of germination and are thus very appropriate to fully characterize germination (McNair et al., 2012). They can especially assess the effects of quantitative covariates (McNair et al., 2012). Cox models are based on a hazard function that expresses the probability of germination at a time t (hazard ratio) and allow determining how specified factors influence this probability. This function takes the following form:

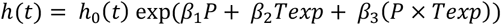

where *P* is the population name (qualitative variable); *Texp* is the experiment temperature; *β*_*i*_ are the coefficients for the base of the exponential factor and exp(*β*_*i*_) represents the hazard ratios; *h*_0_(*t*) the baseline hazards and *t* the germination over time.

The main assumption of Cox proportional hazards models is that the hazard curves for each population and temperature group should be proportional and cannot cross. This assumption was met for experiment temperature, acorn mass, but not for population (Supporting Information: Table S2). To address this issue, we used robust standard errors, which adjust the variance–covariance matrix of the estimated coefficients to remain valid even if the proportional hazards assumption is violated (Lin and Wei, 1989). Model performance was assessed by comparing germination probability curves over time from the Cox model with germination curves from raw data, and by looking at the Nagelkerke R_2_.

We used the survival 3.2-13 R package (Therneau, 2023; Therneau and Grambsch, 2000).

#### 2.5.2. Generalized linear mixed-models of the effects of population and sowing temperature on germination

We fitted linear mixed-effects models to test the effects of the seeds’ climate of origin, experiment temperature (climatic chambers), acorn mass and their interactions on the individual traits G and T0, and on the GS index. We recorded storage duration (time between collection and sowing, up to 22 days) and also tested its effect on traits. We performed stepwise selection on the models to reduce the number of variables and improve their fit, guided by the Akaike Information Criterion (AIC) (function *dredge* of MuMIn 1.47.5 R package (Bartoń, 2023). Storage duration was not significant and excluded from the final models. Acorn mass was also excluded from the final models because its effect was not stable across model stability tests, and we wanted to ensure robust inference.

The models are described below for each germination trait. The choice of the model type depended on the statistical distribution of each trait. In every equation, *ijk* stands for the *i*th acorn of the *j*th population at the *k*th temperature. *Texp* is the experiment temperature; *α*_*n*_ are the coefficients to be estimated; *M* is the mother tree as random effect; *CMD* the Climatic Moisture Deficit; *RMTh* is the Rivas-Martínez Thermicity Index; *TSp* is the maximum spring temperature; *TSn* is the temperature seasonality; *ε* the residuals.

To analyze germination status (G), we built a logistic mixed-effect model using a natural logit link function (function *glmer* from the lme4 1.1-35.1 R package (Bates et al., 2015)). The corresponding model is written as following:

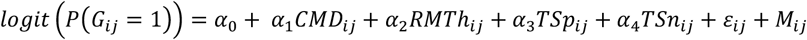

where *G* is the germination status (*G* = 1 means germinated; *G* = 0 means not germinated).

For individual germination time (T0), we built a zero inflated negative binomial mixed-effects model using a log link function (function *glmmTMB* from the glmmTMB 1.1.8 R package (Brooks et al., 2017)) because many acorns germinated very early (0–2 days after sowing), resulting in a large number of low counts. The corresponding model took the following general shape:

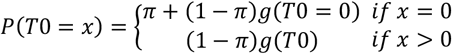

where *T*0 is individual germination time; *π* is the logistic link function written as 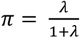 with *λ* the logistic component including regressor variables; *g*(*T*0) is the negative binomial distribution written as 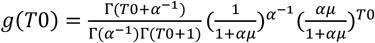 with *μ* the negative binomial component including regressor variables.

Regressor variables are written as follows in the corresponding model:

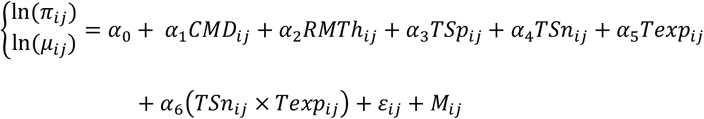

For germination synchrony (GS), we fitted a beta regression model using a logit link function (function *glmmTMB* from the glmmTMB 1.1.8 R package (Brooks et al., 2017)). The general form of the two corresponding models is written as follows:

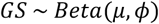

where *GS* is Germination Synchrony; *μ* is the mean parameter (function of predictor variables), modeled by the link function 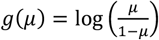 that includes regressor variables; *ϕ* is the precision parameter (determines the dispersion of the beta distribution).

No random effect is needed since Germination synchrony is aggregated by population, mother tree and chamber. Regressor variables are written as follows in the corresponding model:

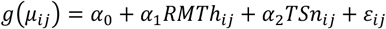

A table of the germination traits, their grouping level, in which models they were used and why, is available at the end of Methods (Table XX).

#### 2.5.3. Models evaluation and multiple comparisons

The variance explained by the models (excluding Cox proportional hazards models; see part 2.5.1) was assessed using different R^2^ metrics because variance partitioning depends on the model type: marginal and conditional for all mixed-effects models, and the ordinary coefficient of determination R^2^ for beta regression model. These calculations were performed using the *model_performance* function from performance 0.10.8 R package (Lüdecke et al., 2021). The generalizability of models was evaluated through cross-validation, using the mean Pearson’s R coefficient, with 0.42% of the data for calibration and 0.58% for validation, repeated 10 times. Residuals plots and Variance Inflation Factors were used to validate the suitability of the models. Tables and plots related to model evaluation are available in Supporting Information (Table S2, Table S3, Fig. S2). All explanatory variables were standardized before running the models.

For the post hoc multiple comparisons we performed between populations and chambers for germination dynamics and germination time rate and synchrony, we used emmeans 1.8.9 R package (Lenth et al., 2024) and multcomp 1.4-25 (Hothorn et al., 2008).

## 3. Results

### 3.1. Climatic characterization of populations

Our populations showed a remarkable north-south climatic gradient. TUN_Tab, SPA_LoAl and TUN_Fer were the warmest populations. TUN_Fer had higher temperature seasonality and was exposed to more water stress than TUN_Tab and SPA_LoAl (Fig. 2). POR_QudSe, SPA_Fue and ITA_Tus had mild conditions. SPA_Fue and ITA_Tus were exposed to higher temperature seasonality and water stress than POR_QudSe (Fig. 2), most likely because the latter is next to the Atlantic Ocean (thus rather exposed to a mild oceanic climate). FRA_MoeMa and FRA_Pad were the coldest populations. FRA_Pad was exposed to higher temperature seasonality and water stress (Fig. 2), most likely because it is located further inland than FRA_MoeMa.

**Fig. 2:**
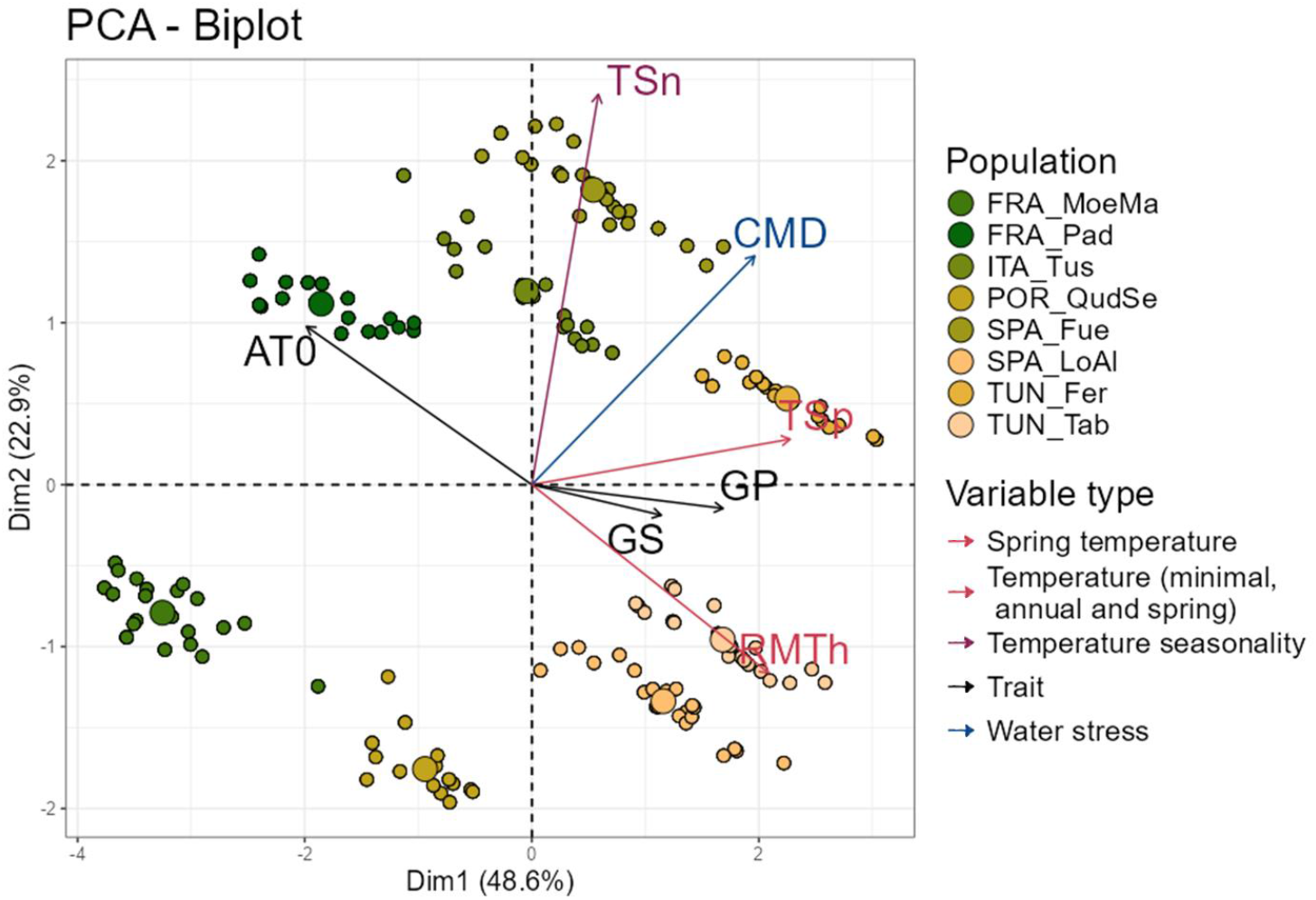
Principal Component Analysis (PCA) representing seeds according to their population’s historical climate (1901-1960 climate variables) and germination performance (indices averaged by population, mother tree and experiment temperature: GP = Germination Percentage; AT0 = Averaged germination time and GS = Germination Synchrony). Small dots represent individual seed sets, each corresponding to a specific population, mother tree, and experiment temperature condition; large dots indicate population-level averages; arrows represent germination traits and population climatic variables, showing their contributions to PCA axes. Clusters of points reflect similar germination/climate behavior, revealing how populations are structured based on their climate and germination responses. The climate variables used are temperature-related factors (TSp = maximum Spring Temperature; TSn = Temperature Seasonality; RMTh = Rivas-Martínez Thermicity Index) and water availability (CMD = Climatic Moisture Deficit). Population colors indicate their origin temperatures (warm colors for warmer populations and cold colors for colder populations, according to RMTh).

### 3.2. Effect of seed origin and temperature on germination dynamics based on Cox proportional hazards model

Cox proportional hazards model showed significant effects of experiment temperature and population on germination probability over time (Table 3).

**Table 3:** Analysis of variance’s results for models of Germination G(0,1) (logistic mixed-effects model), Germination Time T0 (zero inflated negative binomial mixed-effects model), Germination Synchrony GS (beta regression model) and Germination Probability over time (Cox Models). CMD = Climatic Moisture Deficit; TSp = maximum Spring Temperature; TSn = Temperature Seasonality; RMTh = Rivas-Martínez Thermicity Index; Texp = experiment Temperature; Chisq = value of the Chi-squared test; Df = degree of freedom; p = p value. Significant p values (p < 0.05) are shown in bold.

| Covariate | Chisq | Df | p |
| --- | --- | --- | --- |
| Germination probability over time (Cox proportional hazards model) |  |  |  |
| **Population** | **459.73** | **7** | **<0.001** |
| **Texp** | **72.98** | **2** | **<0.001** |
| **Population x Texp** | **27.13** | **14** | **0.018** |
| G (logistic mixed-effects model) |  |  |  |
| CMD | 1 | 1 | 0.317 |
| TSp | 0.3 | 1 | 0.586 |
| TSn | 2.73 | 1 | 0.099 |
| **RMTh** | **11.67** | **1** | **<0.001** |
| T0 (zero inflated negative binomial mixed-effects model) |  |  |  |
| **CMD** | **5.92** | **1** | **0.015** |
| **Texp** | **81.58** | **1** | **<0.001** |
| **RMTh** | **4.31** | **1** | **0.038** |
| TSp | 3.19 | 1 | 0.074 |
| **TSn** | **8.4** | **1** | **0.004** |
| **Texp x TSn** | **6.13** | **1** | **0.013** |
| GS (beta regression model) |  |  |  |
| **RMTh** | **9.27** | **1** | **0.002** |
| TSn | 1.7 | 1 | 0.192 |

Germination probability over time (i.e., the instantaneous probability of germination at time t) increased with seed origin temperature and its climatic moisture deficit and decreased with the seed origin temperature seasonality (Fig. 3, Table S4). On average, seed origin temperature-related variables were able to explain around 36% of the variance. Populations with higher Rivas-Martínez Thermicity index germinate faster (i.e. the warmer half reaching 50% germination 13 days earlier on average than the colder half) while those with lower indices take longer (Table S5), except for TUN_Fer which reach 50% germination before TUN_Tab and SPA_LoAl, most likely because TUN_Fer is drier (Fig. 2) (variance explained by Climatic Moisture Deficit was 17%).

**Fig. 3:**
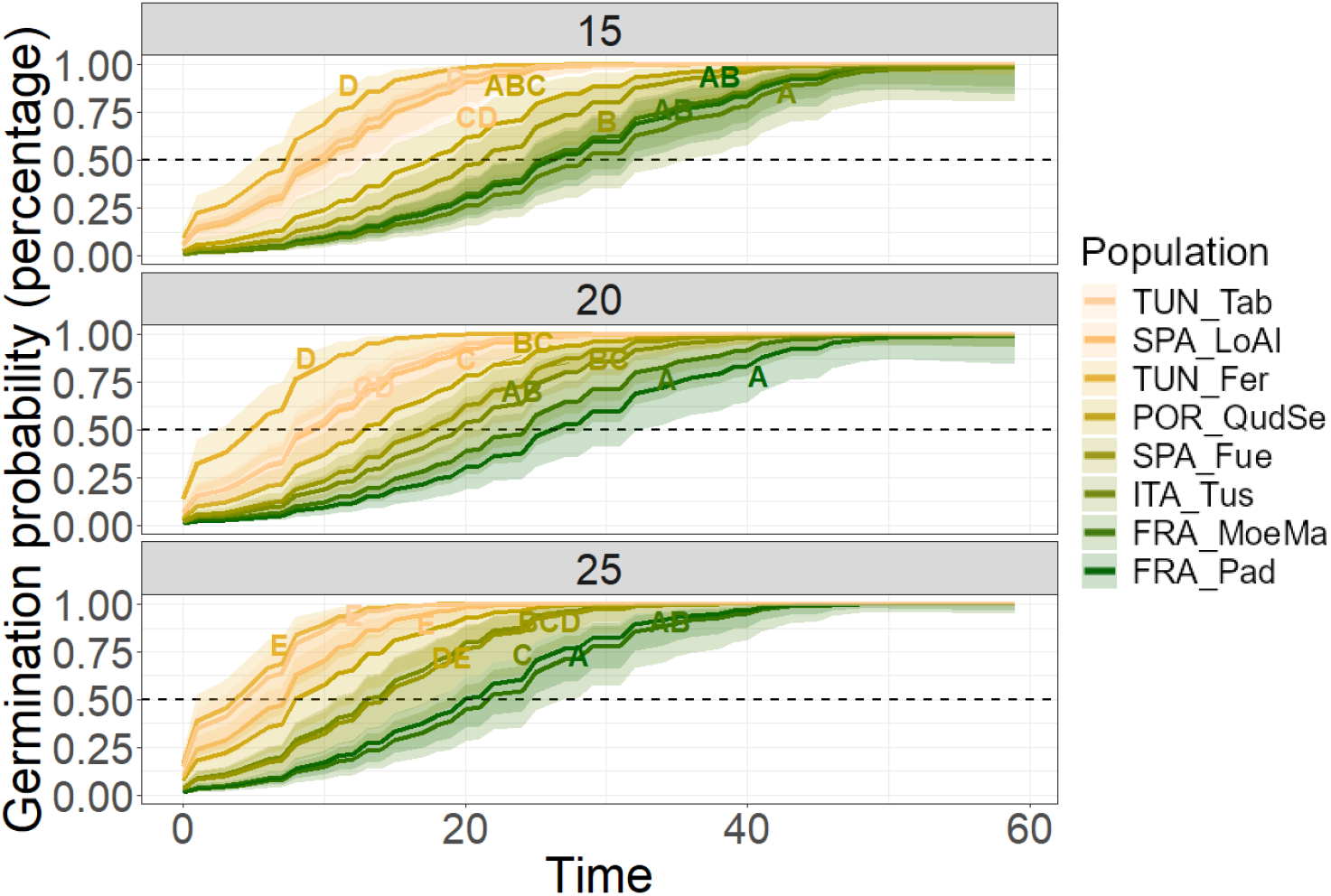
Germination probability over time (days since sowing) predicted by the Cox proportional hazards models at 15°C, 20°C and 25°C and for each population. Populations’ colors indicate their origin temperatures (warm colors for warmer populations and cold colors for colder populations, according to RMTh). Dashed lines indicate the 50% probability of germination and lighter colors the interval of confidence at 5%. Common letters indicate that means do not differ significantly.

The experiment temperature increased germination probability over time (12 ± 7 days at 25°C to reach 50% germination vs. 19 ± 9 days at 15°C) (Fig. S3, Table S5) and its effect was higher on some populations (Fig. S3); seeds from ITA_Tus took 28.4 days at 15°C and 12.9 days at 25 °C (Table S5) to reach 50% germination, i.e. a difference of 15.5 days (Fig. S3). The difference in germination time for POR_QudSe and SPA_Fue was 9.3 and 7 days respectively (Fig. S3). TUN_Fer (3.6 days) and FRA_MoeMa (3.7 days) did not have significant differences in germination probability over time across temperatures (Fig. S3). The Nagelkerke R^2^ of the model was 0.42 (Table S3).

### 3.3. Effect of seed origin and temperature on germination percentage, timing and synchrony based on mixed-effects models

#### 3.3.1. Germination percentage

Only population climatic variables had a significant effect on germination percentage (Table 3). Germination percentage increased along with population temperature and temperature seasonality (Fig. 4, Table S4), with warmer populations germinating more than colder ones (TUN_Fer (97 ± 7 %), TUN_Tab (94 ± 9 %), SPA_LoAl (95 ± 10 %) and ITA_Tus (96 ± 9 %) vs. SPA_Fue (83 ± 20 %), POR_QudSe (77 ± 30 %), FRA_MoeMa (68 ± 20 %) and FRA_Pad; (66 ± 30 %) (Fig. 4, Fig. S4).

**Fig. 4:**
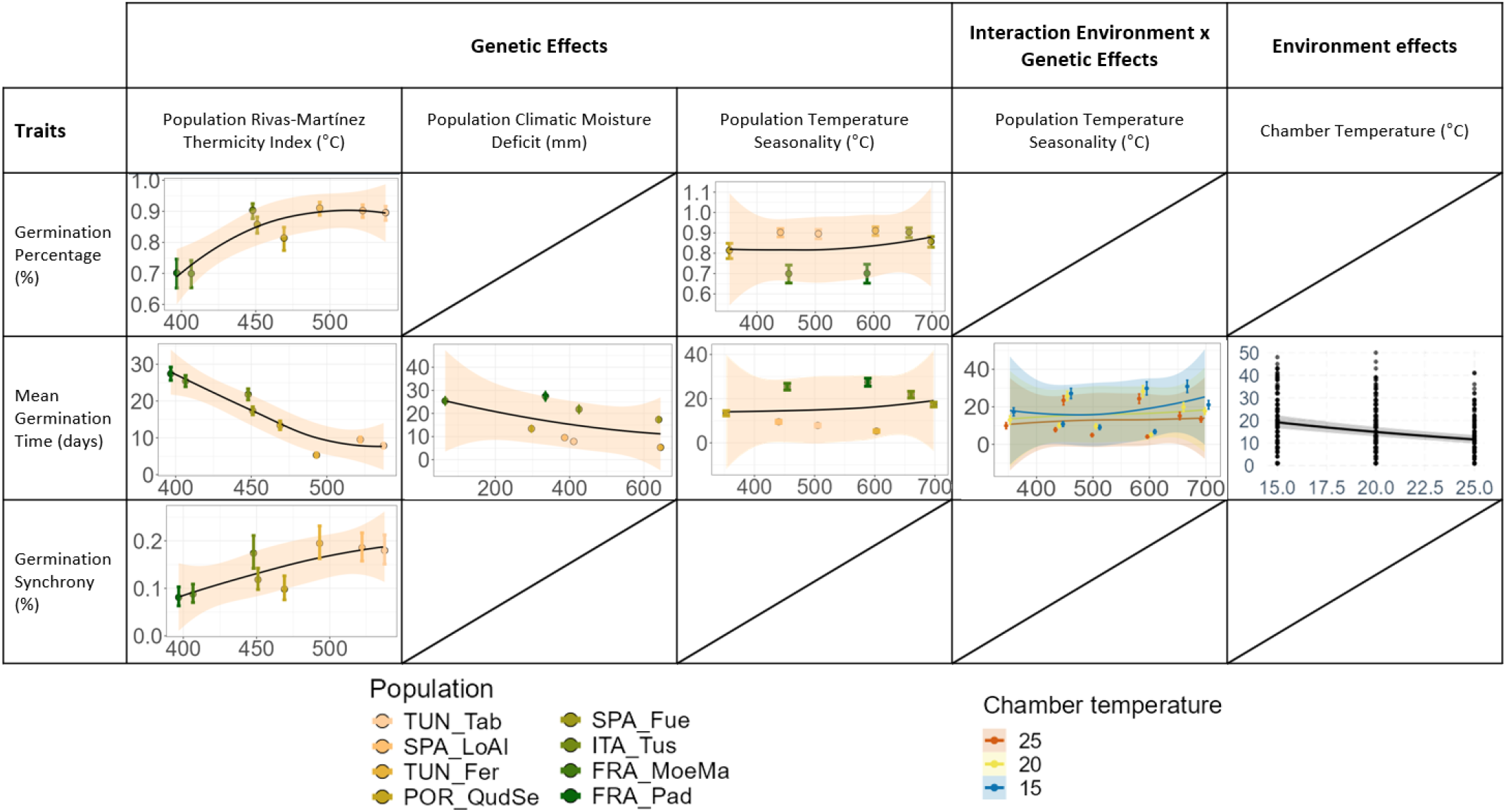
Summary of the population effects (i.e. population climatic variables), environment effects (i.e. experiment temperature), and their interaction (i.e. experiment temperature effect according to population) on germination metrics. Non-significant effects are represented by crossed-out boxes. The effect of experiment temperature on germination time was visualized using model-predicted values, averaged across populations (i.e., marginal means) where this effect was significant, to obtain three curves representing the three experiment temperatures along the gradient of population temperature seasonality. The magnitude of germination time plasticity is represented by the difference between germination time curves at 15°C (blue), 20°C (yellow) and 25°C (red). Populations’ colors indicate their origin temperatures (warm colors for warmer populations and cold ones for colder populations).

Pearson Correlation Coefficient of the model was 0.82. The conditional R^2^ was 0.32 and the marginal R^2^ was 0.21. Additional performance metrics like AIC and RMSE are included in Table S3.

#### 3.3.2. Germination time

Experiment temperature, seed origin temperature, and their interaction had a significant effect on germination time (Table 3). Germination time decreased along with rising temperatures (time to germinate was 12.2 ± 9.3 days in the 25°C chamber vs. 15.9 ± 10.7 days in the 20°C chamber vs. 18.4 ± 12.5 days in the 15°C chamber) (Fig. 4 and Table S4); acorns exposed to higher temperatures germinated earlier than acorns at lower temperatures.

Germination time decreased with increasing seed origin temperature and water stress and increased with increasing temperature seasonality (Fig. 4 and Table S4). TUN_Fer was the first population to germinate (5.3 ± 3.0 days), and FRA_MoeMa (25.5 ± 6.0 days) and FRA_Pad (27.2 ± 3.9 days) were the last ones (Fig. 4, Fig. S4).

The interaction between seed origin temperature seasonality and experiment temperature was significant (Table 3). Experiment temperature had a higher effect on seeds with increasing temperature seasonality, particularly continental populations as ITA_Tus where germination time was on average doubled in the coldest chamber compared to the warmest chamber (+ 15.7 days), and to a lesser extent for SPA_Fue (+ 7.8 days) and POR_QudSe (+ 7.4 days) (Fig. 4, Table S6). The interaction effect was smaller for SPA_LoAl (+ 3.0 days) and TUN_Tab (+ 4.1 days) (Fig. 4, Fig. S5, S6, Table S6). In contrast, TUN_Fer, FRA_MoeMa and FRA_Pad did not have significant differences in mean germination time across temperatures (Fig. 4, Fig. S5, S6). Pearson Correlation Coefficient of the model was 0.68. The conditional R^2^ was 0.60 and the marginal R^2^ was 0.44 (Table S3).

#### 3.3.3. Germination synchrony

Only seed origin temperature had a significant effect on germination synchrony (Table 3). Germination synchrony increased with seed origin temperature (Fig. 4, Table S4). Seeds from FRA_Pad showed the lowest germination synchrony (FRA_Pad: 0.08 ± 0.02 %; Other populations’ average: 0.16 ± 0.04 %) (Fig. 4, Fig. S4). Pearson Correlation Coefficient of the model was 0.35. The R^2^ was 0.13 (Table S3).

## 4. Discussion

The genetic clines found in cork oak germination percentage, time and synchrony were mostly related to population temperature, as already observed for traits measured in adult trees where adaptation to temperature is prevalent, e.g. height growth, budburst (Saxe et al., 2001). Surprisingly, the low germination synchrony of cork oak from colder populations suggests a bet-hedging adaptive strategy of these populations to unpredictable frost events that would impair seedlings’ survival. Germination timing was the only trait that varied depending on the experiment temperature. This variation occurred primarily in acorns originating from populations with higher temperature seasonality, suggesting low overall plasticity in germination timing in response to the experiment temperature.

### 4.1. Germination percentage, time and synchrony are determined by climate at seed origin

#### 4.1.1. Germination percentage increases with seed origin temperature

Germination percentage was mostly determined by seed origin temperature, with seeds from warmer populations being more likely to germinate than those from colder ones, regardless of the sowing temperature (Fig. 4). Similar results have been reported for cork oak with partially desiccated acorns (Benito Garzón et al., 2024), other Mediterranean oaks as *Quercus ilex* (García-Nogales et al., 2016), and for temperate species with recalcitrant seeds like *Acer pseudoplatanus* and *Aesculus hypocastanum* (Daws et al., 2006, 2004). These findings suggest that, in species with recalcitrant seeds, germination percentage increases along with temperature at seed origin. This could be related to the physiology of recalcitrant seed development. Indeed, heat sum during this period is usually related to an enhanced vigor, maturation status and germinability (Daws et al., 2006, 2004; Fernández-Pascual et al., 2019). Thus, warmer conditions during seed development in southern populations, may have increased acorn viability and germinability compared to northern, colder populations.

In addition, the higher germination of warm-origin seeds indicates local adaptation to warm conditions, where maintaining higher germinability may confer a fitness advantage. In cork oak, this pattern might be related to its evolutionary history. *Quercus suber* and other species from the *cerris* section originated during the Miocene when Mediterranean basin climate shifted from subtropical to more seasonal and warm-temperate (Denk et al., 2023), implying adaptation to conditions warmer and wetter than today.

#### 4.1.2. Seeds from warmer and drier origins germinate earlier than those from other origins

Our results showed a clear cline in germination time: the warmer and drier the seed origin, the earlier germination occurred (Fig. 4). Similar population-level effects on germination time have been reported for cork oak and other Mediterranean trees (Benito Garzón et al., 2024; Vicente and Benito Garzón, 2024), as well as in other plant species (e.g. Meyer et al., 1989; Zettlemoyer et al., 2017).

Earlier germination in warm and dry environments provides two advantages: (i) it reduces the period of desiccation risk between acorn shedding and germination (Joët et al., 2016, 2013), and (ii) it provides seedlings a longer growth time window before the onset of summer drought. This is consistent with evidence that southern populations exhibit greater development of morphological traits related to water use, enabling seedlings to establish before the onset of summer drought (Ramírez-Valiente et al., 2022; Vicente et al., 2025). Conversely, delayed germination in seeds from cold and wet origins may represent an adaptive strategy to avoid late frost damage (Muffler et al., 2016).

Our findings support that germination time, like other phenological traits such as budburst and flowering timing, is highly adapted to local conditions, reflecting strong selection by temperature and water availability (Augspurger, 2008; Billington and Pelham, 1991; Chmura, 2006).

#### 4.1.3. Seeds from cold origins show the lowest synchrony

Germination synchrony is influenced by seed dormancy. Dormant seeds typically spread germination over longer time periods than non-dormant seeds (Yang et al., 2013); therefore, most recalcitrant seeds (being largely non-dormant) are expected to germinate with high synchrony to avoid desiccation after shedding (Maleki et al., 2023). However, our results with cork oak indicate that synchrony also varies with temperature at seed origin, suggesting an adaptive response. Seeds from the coldest populations (FRA_Pad, FRA_MoeMa) showed lower synchrony, likely as an adaptive response to spread the exposure to frost events. A similar pattern has been observed in red oak (*Quercus pagoda*) in the southeastern United States, where higher germination temperatures were associated with higher synchrony (Hawkins, 2019).

These findings suggest that even in recalcitrant species, germination synchrony decreases under less favourable or less predictable conditions, paralleling the ecological strategies of orthodox species. In such environments, reduced synchrony functions as a bet-hedging mechanism that spreads risk across time. For instance, in alpine habitats it reduces the risks of frost damage in early-spring (Fernández-Pascual et al., 2021), whilst in deserts and arid environments, it helps to cope with the precipitation fluctuations, contributing also to lessen competition (Bhatt et al., 2020; Gremer et al., 2016; Nimac et al., 2018).

### 4.2. Sowing environment only affects germination time

The lack of variation in germination percentage across the three experimental temperatures suggest low plasticity to temperature in this trait (Fig. 4), as already observed for other Mediterranean species (Vicente and Benito Garzón, 2024). By contrast, in a study with cork oak using similar experimental temperatures, germination percentage dropped sharply when acorns were exposed to desiccation before sowing, and germination varied greatly between temperature treatments (Benito Garzón et al., 2024). In our study, however, germination percentages were nearly identical across temperatures (69%, 74%, and 73%, at 15°C, 20°C, and 25°C), suggesting that desiccation sensitivity may condition germination response to temperature variation.

Similarly, germination synchrony did not vary with experimental temperature (Fig. 4). This was unexpected, given that previous studies found a significant response of this trait to temperature variation (Bhatt et al., 2020; Fernández-Pascual et al., 2021; Nimac et al., 2018). However, these studies were all focused on species with orthodox seeds. Nonetheless, it cannot be ruled out that the responses observed in germination percentage and synchrony might become more pronounced across a broader temperature range. Experiments conducted under colder or warmer regimes may yield different results.

In contrast, germination time did respond to sowing temperatures: seeds germinated earlier under warmer conditions, as occurred in other studies showing that rising temperatures advance germination time (Yuan et al., 2023). Under future warming conditions, this may translate into earlier germination across cork oak distribution range. In addition, it is worth noting that the effect of increasing temperatures hastening germination was particularly relevant in the acorns originating from Tuscania (Italy, ITA_Tus population). This could be related to the temperature seasonality of this provenance, which is the highest among our sampled populations. Indeed, trees from populations exposed to greater climatic variability are usually reported to display greater plasticity in their functional traits (Janzen, 1967; Lázaro-Nogal et al., 2015; Pratt and Mooney, 2013). In this context, greater plasticity could confer an adaptive advantage, as it may allow faster adjustments to manage unpredictable environments (Valladares et al., 2014). Thus, trees from this population, with recurrent harsher winters and summers, may have evolved to increase germination time plasticity as an adaptive strategy to increase the chances for germination.

### 4.3. Limitations and perspectives

Our experiment was not designed to assess the effects of water availability on germination, and we lacked the driest cork oak populations in which germination and its plasticity generally increase (Vicente and Benito Garzón, 2024). Hence, caution is needed when interpreting our results, particularly in the context of climate change where drought is expected to increase. Furthermore, we used acorns coming from a single year fructification, hence the inter-annual variability of acorn production and quality usually occurring in most *Quercus* species (Mechergui et al., 2023) is not included.

The strong effect of seed origin on percentage, time and synchrony of germination suggests that these traits should be taken into account in climate resilience strategies including assisted migration programs, which are generally based on adult traits performance rather than that of early stages (Chakraborty et al., 2024). The adaptive response found for germination synchrony was unexpected and further investigation on recalcitrant seeds germination dynamics in response to increased temperatures is needed to confirm it. Finally, the integration of germination studies in a wider regeneration context is needed to fully understand the chances of trees to survive under warmer and drier conditions.

## Supporting information

Electronic Supplementary material

## Conflict of Interest Statement

We declare that the authors have no competing interests as defined by Springer, or other interests that might be perceived to influence the results and/or discussion reported in this paper.

## Contributions by the Authors

MBG and EV designed the study. EV, MC and MBG conducted the reciprocal sowing experiment and monitored germination. MC analyzed the data. MC and MBG wrote the manuscript. All the authors collected the acorns and edited the final manuscript. MBG acquired the financial support needed for this project.

## Funding

This research has been supported by the Nouvelle Aquitaine Regional project SUBER. MC was funded by a PhD doctoral grant from ECODIV and the Nouvelle Aquitaine Regional project SUBER. ED was funded by EU-funded project SUPERB Green Deal H2020. NVP was funded by IJC2020-044557-I/ AEI / 10.13039/501100011033.

## Notes

### Competing Interest Statement

The authors have declared no competing interest.

