## Supplementary material for "Seed origin determines cork oak germination: the warmer the higher, faster and more synchronized": Electronic Supplementary material

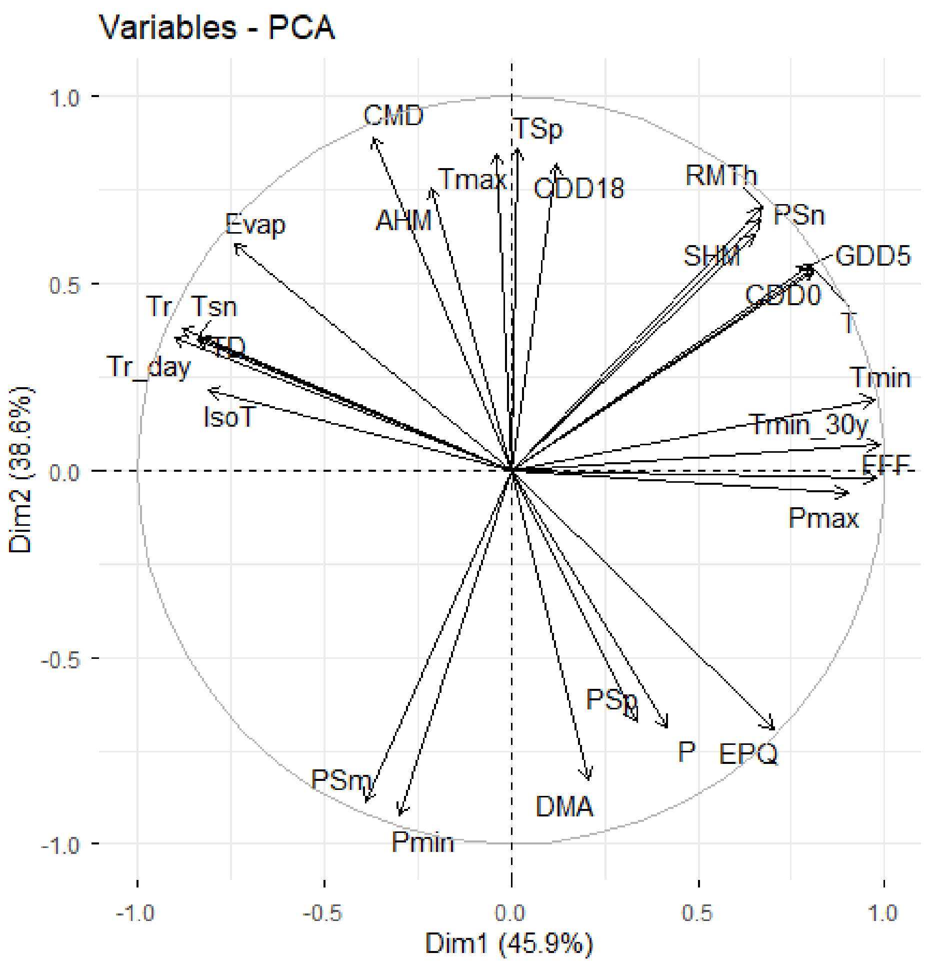

**Fig. S1:** Principal Component Analysis (PCA) correlation circle of all climatic variables in order to select non-correlated variables. All variables were retrieved from (ClimateDT; https://www.ibbr.cnr.it/climate-dt/ Marchi et al., 2024). TSp = maximum spring temperature; PSp = mean spring precipitation; T = mean annual temperature; Tr_day = mean diurnal temperature range; IsoT = isothermality; Tsn = temperature seasonality; Tr = annual temperature range; P = mean annual precipitation; Pmax = precipitation of wettest month; Pmin = precipitation of driest month; PSn = precipitation seasonality; Evap = monthly standardised potential evapotranspiration index; CMD = climatic moisture deficit; Tmin = minimum temperature of coldest month; PSm = mean summer precipitation; Tmax = maximum temperature of warmest month; CDD0 = chilling degree-days - above 0 °C; GDD5 = growing degree-days - above 5 °C; CDD18 = cooling degree-days - above 18 °C; FFF = frost-free days per year; DMA = De Martonne aridity index; EPQ = Emberger pluviometric quotient; RMTh = Rivas-Martínez thermicity index; AHM = annual heat-moisture index; SHM = summer heat-moisture index; TD = continentality index (temperature difference between warmest & coldest months); Tmin_30y = minimum temperature averaged over the last 30 years.

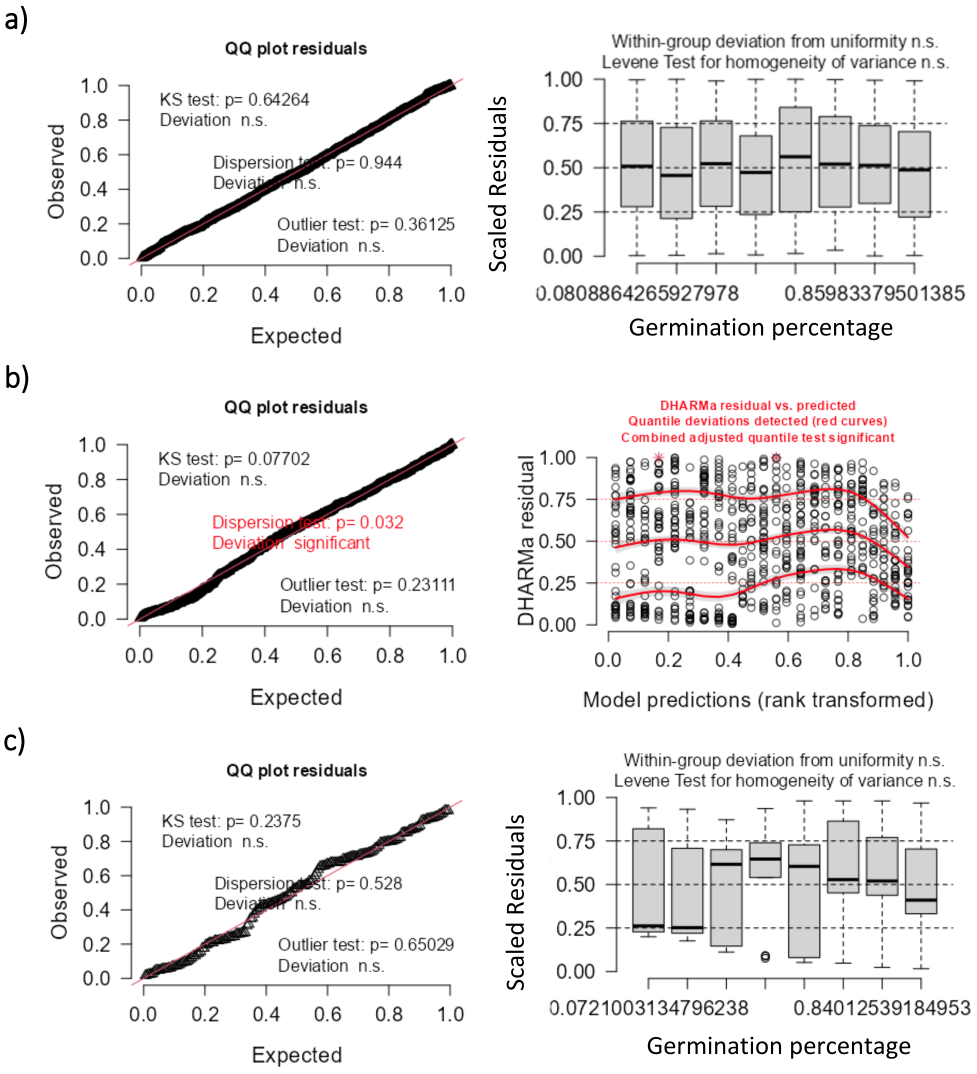

**Fig. S2:** Plots of residuals from regression models. Germination (G) model in a); Germination time (T0) model in b); Germination synchrony (GS) model c).

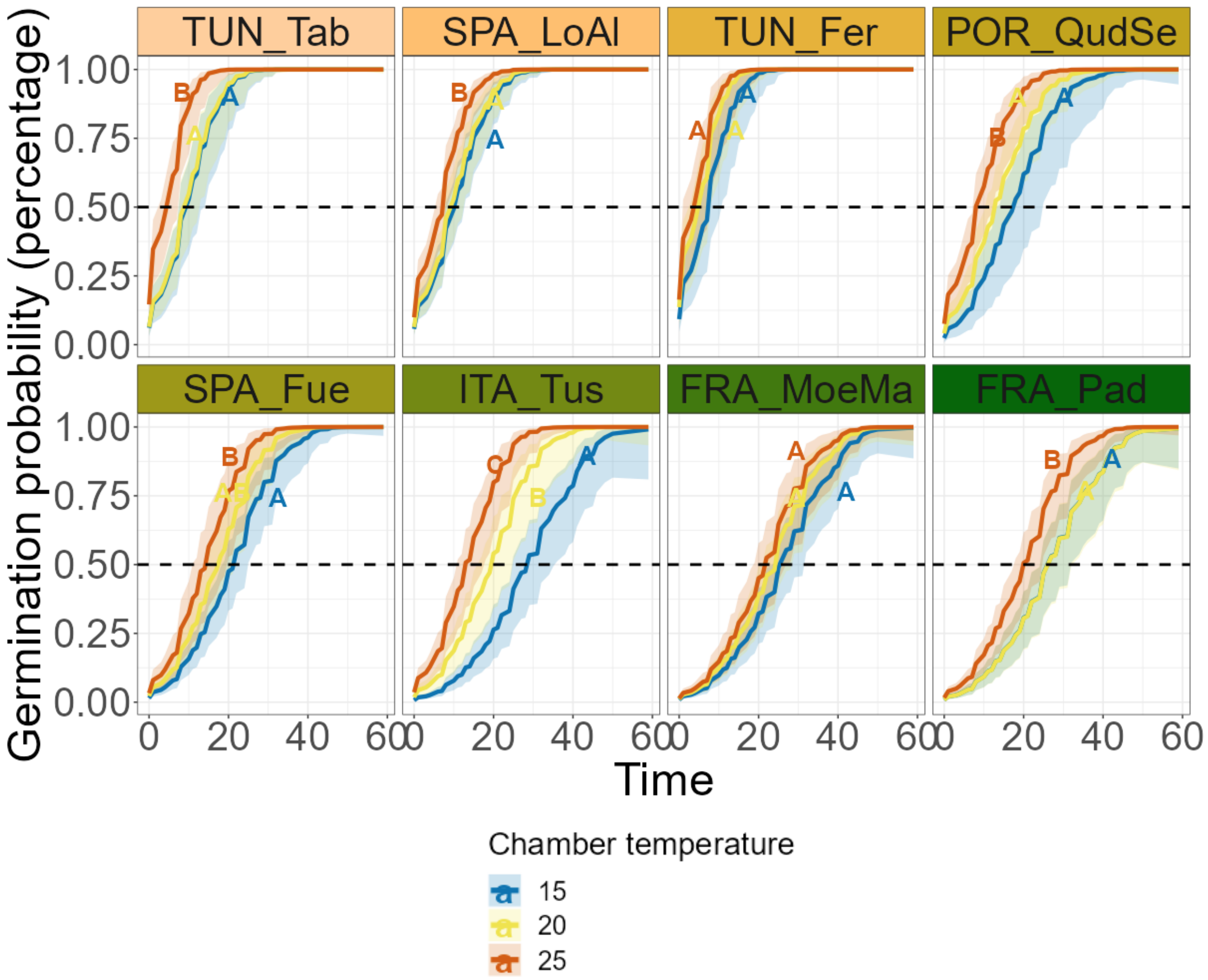

**Fig. S3:** Germination probability over time (days since sowing) estimated by the Cox proportional hazards models at 15°C, 20°C and 25°C and for each population. Populations' color indicates their origin temperatures (warm colors for warmer populations and cold colors for colder populations, according to RMTh). Differences between experiment temperatures are enlightened, with populations ordered by temperature seasonality. The dashed line indicates the 50% probability of germination and the lighter colors the interval of confidence at 5%. Common letters indicate that means do not differ significantly.

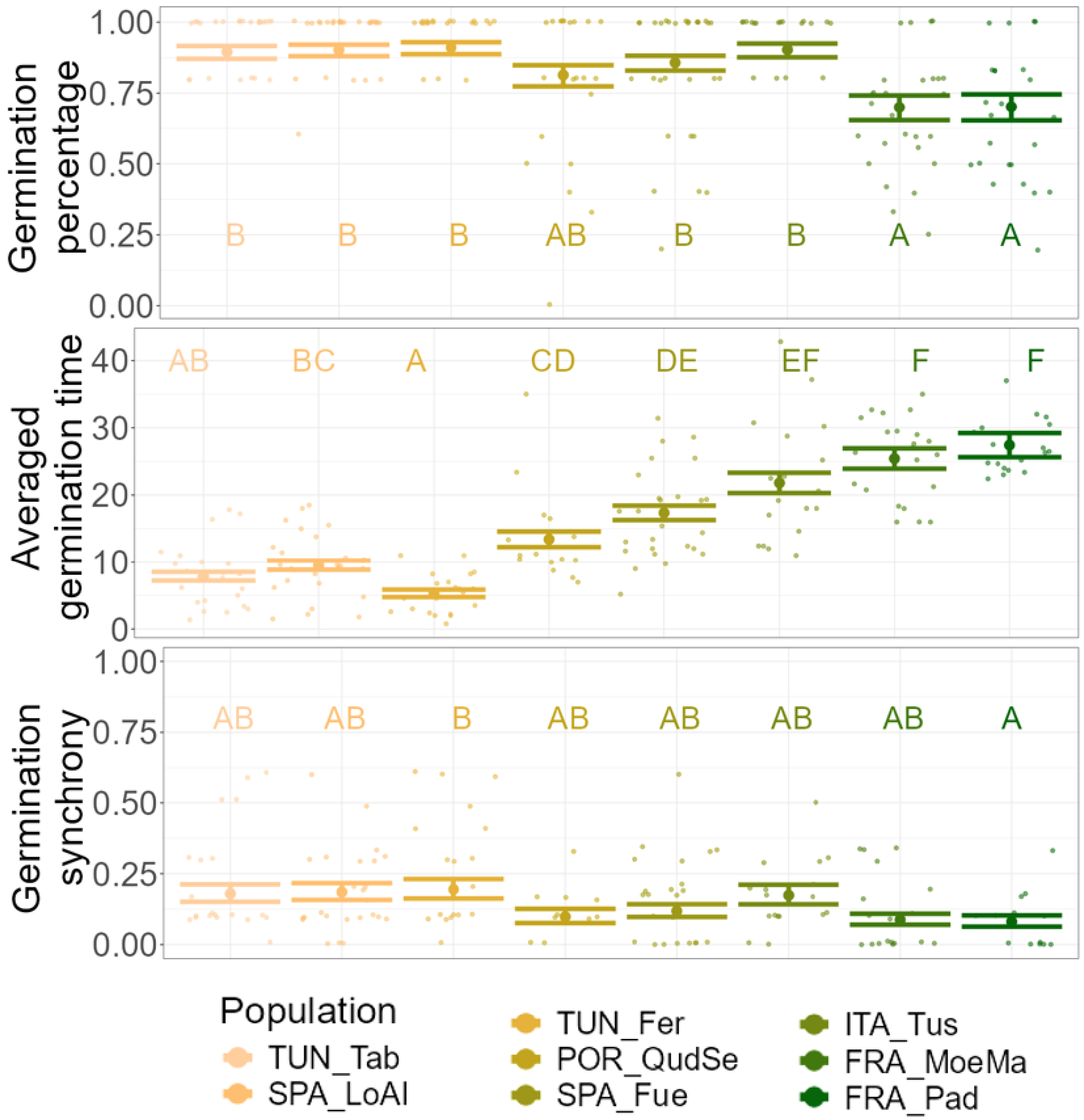

**Fig. S4:** Mean and standard deviation of germination percentage (GP), Averaged germination time (AT0), and germination synchrony from pairwise comparisons between populations. Common letters indicate that means do not differ significantly. Semi-transparent points are the raw values. Populations' color indicates their origin temperatures (warm colors for warmer populations and cold colors for colder populations, according to RMTh).

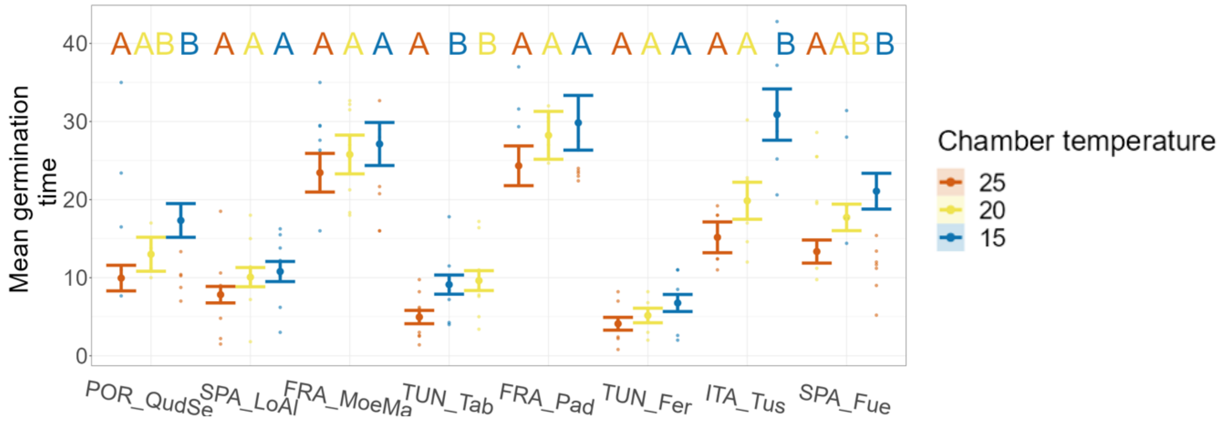

**Fig. S5:** Mean and standard deviation of germination time from pairwise comparisons between populations and experiment temperatures, with populations ordered by temperature seasonality. Common letters indicate that means do not differ significantly. Transparent points are the raw values.

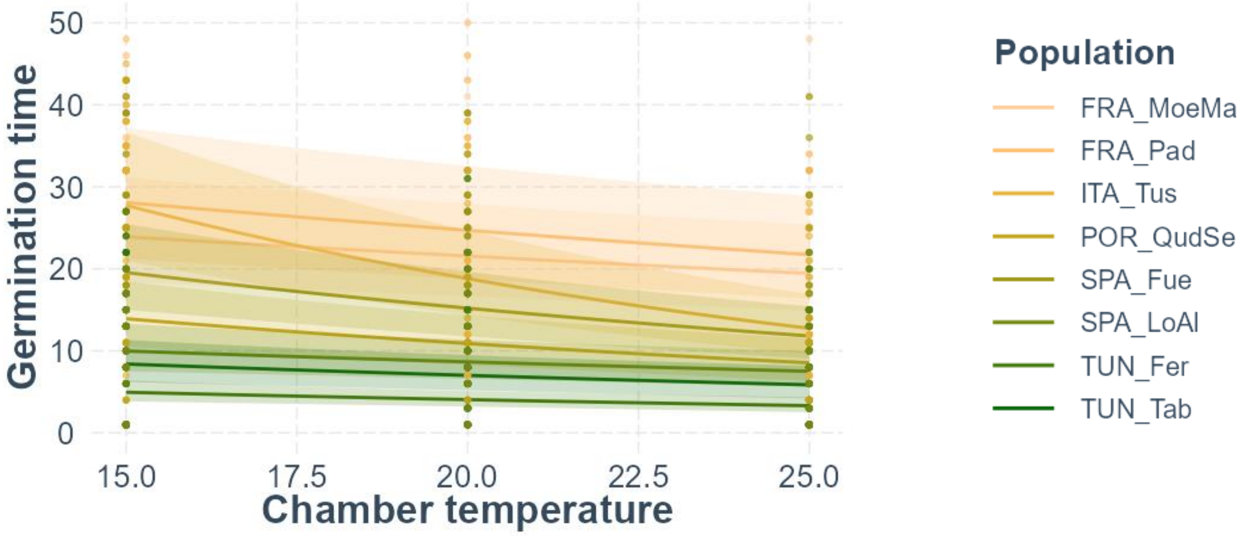

**Fig. S6:** Interaction between experiment temperature effect and population, calculated with the germination time mixed-effects model. Populations' color indicates their origin temperatures (warm colors for warmer populations and cold colors for colder populations, according to RMTh).

**Table S1:** Mean maximum spring temperatures (TSp), mean winter temperatures (TWt), and mean autumn temperatures (TAt) for each population, averaged over the period 1901–1960. All temperature variables were retrieved from (ClimateDT; https://www.ibbr.cnr.it/climate-dt/ Marchi et al., 2024). Populations are sorted by Rivas-Martínez Thermicity Index.

| Population | TSp | TWt | TAt |
|---|---|---|---|
| **TUN_Tab** | 17,99 | 13,67 | 21,90 |
| **SPA_LoAl** | 17,60 | 13,97 | 20,31 |
| **TUN_Fer** | 18,13 | 12,18 | 21,63 |
| **POR_QudSe** | 15,73 | 12,92 | 18,54 |
| **SPA_Fue** | 16,82 | 9,65 | 19,06 |
| **ITA_Tus** | 16,92 | 9,95 | 19,02 |
| **FRA_MoeMa** | 14,82 | 10,50 | 17,33 |
| **FRA_Pad** | 16,72 | 9,42 | 17,95 |

**Table S2:** Proportional Hazards assumption in the Cox models. Non-significant p values (p > 0.05) are shown in bold, to highlight the variables for which the Proportional Hazards assumption is met. Chisq the value of the Chi-squared test; Df the degree of freedom; p the p value; CMD the Climatic Moisture Deficit; TSp the maximum Spring Temperature; TSn the Temperature Seasonality; RMTh the Rivas-Martínez Thermicity Index; Texp the experiment Temperature.

| Covariate | chisq | df | p |
|---|---|---|---|
| Population | 60.59 | 7 | <0.001 |
| **Texp** | **0.097** | **2** | **0.953** |

| | | | |
|---|---|---|---|
| Population x Texp | 28.97 | 14 | 0.011 |
| GLOBAL | 74.24 | 23 | <0.001 |

**Table S3:** Models performance measures. G: Germination Status; T0: Germination Time; GS: Germination Synchrony ; AIC the Akaike Information Criterion (measure of model quality that balances goodness-of-fit and model complexity); AICc the corrected Akaike Information Criterion (adjusted for small sample sizes); BIC the Bayesian Information Criterion (penalizes model complexity more strongly); $R^2$ cond. the conditional R-squared (proportion of variance explained by both fixed and random effects); $R^2$ marg. the marginal R-squared (proportion of variance explained by fixed effects); $R^2$ the R-squared (proportion of variance explained by the model in general); $R^2$ Nagelkerke, the Nagelkerke's R-squared (a pseudo-$R^2$ measure for generalized linear models, scaled between 0 and 1); RMSE the Root Mean Square Error (average magnitude of model prediction errors); R Pearson the Pearson correlation coefficient between observed and predicted values.

| **Model** | **AIC** | **AICc** | **BIC** | **$R^2$ cond.** | **$R^2$ marg.** | **$R^2$** | **$R^2$ Nagelkerke** | **RMSE** | **R Pearson** |
|---|---|---|---|---|---|---|---|---|---|
| **G model** | 765.667 | 765.755 | 794.875 | 0.317 | 0.214 | | | 0.337 | 0.825 |
| **T0 model** | 5331.598 | 5331.876 | 5378.456 | 0.597 | 0.444 | | | 6.819 | 0.68 |
| **GS model** | -151.695 | -151.162 | -142.167 | | | 0.181 | | 0.157 | 0.153 |

| | | | | | | | |
|---|---|---|---|---|---|---|---|
| **Cox model** | 8652.2 | 8653.6 | 8759.9 | | | 0.472 | 1.097 |

**Table S4:** Parameters for models of Germination G (logistic mixed-effects model), Germination Time T0 (zero inflated negative binomial mixed-effects model), and Germination Synchrony GS (beta regression model). Estimate are the coefficients of regression; Std. Error the standard errors of the coefficients; Test value the test (Z-test) values of the models, p the p value. Significant p values ($p < 0.05$) are shown in bold. CMD the Climatic Moisture Deficit; TSp the maximum Spring Temperature; TSn the Temperature Seasonality; RMTh the Rivas-Martínez Thermicity Index; Texp the experiment Temperature.

| Covariate | Estimate | Std. Error | Test value | p |
|---|---|---|---|---|
| G (logistic mixed-effects model) | | | | |
| **(Intercept)** | **2.1** | **0.16** | **12.82** | **<0.001** |
| CMD | -0.33 | 0.33 | -1 | 0.317 |
| TSp | 0.16 | 0.3 | 0.54 | 0.586 |
| TSn | 0.54 | 0.33 | 1.65 | 0.099 |
| **RMTh** | **1.12** | **0.33** | **3.42** | **<0.001** |
| T0 (zero inflated negative binomial mixed-effects model) | | | | |
| **(Intercept)** | **2.61** | **0.04** | **58.9** | **<0.001** |
| **CMD** | **-0.24** | **0.1** | **-2.43** | **0.015** |
| **Texp** | **-0.15** | **0.02** | **-8.58** | **<0.001** |
| **RMTh** | **-0.22** | **0.11** | **-2.08** | **0.038** |

| | | | | |
|---|---|---|---|---|
| TSp | -0.18 | 0.1 | -1.79 | 0.074 |
| **TSn** | **0.3** | **0.11** | **2.84** | **0.004** |
| **Texp x TSn** | **-0.05** | **0.02** | **-2.48** | **0.013** |
| GS (beta regression model) | | | | |
| **(Intercept)** | **-1.88** | **0.14** | **-13.78** | **<0.001** |
| **RMTh** | **0.37** | **0.12** | **3.04** | **0.002** |
| TSn | 0.16 | 0.12 | 1.31 | 0.192 |

**Table S5:** Time (in days) to reach 50% germination (T50) for every population and the three chambers temperature, means and standard deviations across temperatures, and Rivas-Martínez Thermicity Index (RMTh). Populations are sorted by Rivas-Martínez Thermicity Index.

| Population | 15°C | 20°C | 25°C | Mean | Standard deviation | RMTh |
|---|---|---|---|---|---|---|
| **TUN_Tab** | 9.2 | 8.8 | 4.4 | 7.47 | 2.66 | 537 |
| **SPA_LoAl** | 10.1 | 9.2 | 7.2 | 8.83 | 1.48 | 522 |
| **TUN_Fer** | 7.3 | 4.8 | 3.7 | 5.27 | 1.84 | 493 |
| **POR_QudSe** | 17.3 | 12.7 | 8 | 12.67 | 4.65 | 469 |
| **SPA_Fue** | 21.2 | 17.4 | 14.2 | 17.60 | 3.50 | 451 |
| **ITA_Tus** | 28.4 | 19.4 | 12.9 | 20.23 | 7.78 | 448 |
| **FRA_MoeMa** | 25.3 | 24.2 | 21.6 | 23.70 | 1.90 | 407 |
| **FRA_Pad** | 26 | 26.1 | 19.9 | 24.00 | 3.55 | 397 |

**Table S6:** Averaged germination time (AT0, in days) per population for the coldest (15°C) and the warmest (25°C) climatic chambers. Populations are sorted by Rivas-Martínez Thermicity Index. Values are reported only for populations in which the interaction between experimental temperature (Texp) and population temperature seasonality (TSn) was significant. For each population and chamber, the mean (Mean) and standard deviation (SD) of $AT_0$ are shown. Populations for which the interaction was not significant are indicated as such.

| Population \ Texp | 15°C | | 25°C | |
|---|---|---|---|---|
| | **Mean** | **SD** | **Mean** | **SD** |
| **TUN_Tab** | 9.1 | 4.4 | 5.0 | 3.0 |
| **SPA_LoAl** | 10.8 | 4.3 | 7.8 | 5.1 |
| **TUN_Fer** | Interaction Texp and TSn not significant | | | |
| **POR_QudSe** | 17.3 | 10.3 | 9.9 | 2.3 |
| **SPA_Fue** | 21.1 | 6.2 | 13.3 | 5.3 |
| **ITA_Tus** | 30.9 | 8.1 | 15.2 | 3.6 |
| **FRA_MoeMa** | Interaction Texp and TSn not significant | | | |
| **FRA_Pad** | Interaction Texp and TSn not significant | | | |
